# Application of sequence semantic and integrated cellular geography approach to study biogenesis of exonic circular RNA

**DOI:** 10.1101/2021.10.26.465935

**Authors:** Rajnish Kumar, Tapobrata Lahiri, Manoj Kumar Pal

## Abstract

**Background:** Concurrent existence of lncRNA and circular RNA at both nucleus and cytosol within a cell at different proportion is well reported. Secondly, information on genes transcribing both circular and lncRNAs along with total number of RBP binding sites for both of these RNA types is extractable from databases. This study showed how these apparently unconnected pieces of reports could be put together to build a model leading to biogenesis of circular RNA.

**Results:** As a result of this study, a model was built under the premises that, sequences with special semantics were molecular precursors in biogenesis of circular RNA which occurred through catalytic role of some specific RBPs. The model outcome was further strengthened by fulfillment of three logical lemmas which were extracted and assimilated in this work using a novel data analytic approach, Integrated Cellular Geography. Result of the study was found to be in well agreement with proposed model. Furthermore this study also indicated that biogenesis of circular RNA was a post-transcriptional event.

**Conclusions:** Overall, this study provides a novel systems biology based model under the paradigm of Integrated Cellular Geography which can assimilate independently performed experimental results and data published by global researchers on RNA biology to provide important information on biogenesis of exonic circular RNAs considering lncRNAs as precursor molecule.

## Background

A complete physicochemical mechanism of biogenesis of circular RNAs is still under scrutiny. In some study back-splicing supported by the presence of inverted repeats in flanking Introns are considered as root cause of their biogenesis [1,2], while in other study involvement of some trans-factors like RNA Binding Proteins (RBPs) are reported to induce the event of circularization [3–5]. Apart from this, Exon skipping events are also considered to be the influencing factor of the circularization event [6]. Recently, RBPs which are of immune factor types are reported to be involved in Circular RNAs biogenesis [7].

Interestingly, both long non-coding RNAs (lncRNAs) and Circular RNAs with length greater than 200 bp are found to share many common properties [8,9]. Both Circular RNAs and lncRNAs are reported to work as biomarker in case of cancers and other diseases [10]. Circular RNAs differ from mRNA mainly in terms of topology, stability and translational capability and its biogenesis competes with splicing from pre-mRNA [4]. Long non-coding RNAs (lncRNAs) are mRNAs like transcripts which was first reported in Xist gene of mouse [11]. As reported in [12], in context of coding potential, lncRNAs primarily lack Open Reading Frame (ORF). Regarding differences, lncRNAs differ from mRNAs through their larger transcript sizes, longer exon lengths, low conservation of sequence, relatively low expression and more specific expression profile which are considered as features discriminating them from mRNAs [13,14]. lncRNAs can be classified at different levels based on their function, localization and biogenesis[15].

Role of lncRNAs in post-transcriptional regulation is well reported [16]. lncRNAs are reported to be molecular address code particularly in nucleus [17]. Their associations with disease as well as cellular functions are so high that they are considered as multitasking molecules inside the cell [18]. Some of the lncRNAs undergo post-transcriptional processing resulting in alternative forms that are different than other reported lncRNAs [19,20]. From these reports it is noticeable that field of lncRNAs is expanding. Recently a new type of lncRNAs are reported that are truly exonic like mRNAs having both ends closed [21,22]. It is however intriguing to observe that another member of RNA world, Circular RNAs (circRNAs) resembles lncRNAs on many aspects mining of which may provide a pack of information on roles and biogenesis of these biomolecules.

In the present study the process of circularization of RNA was investigated by considering results of research works so far published under an introductory data processing framework, Integrated Data Geography (IDG) in general and Integrated Cellular Geography (ICG) in particular following well known Integrated Geography (IG) approach [23]. Under this framework experimental outputs of published reports were considered as lemmas to utilize them as necessary and sufficient conditions for validation of the model of circular RNA Biogenesis proposed in this work. In the backdrop of many published results available through various investigations, introduction of this approach appeared to be the need of the hour for convergence of such apparently unconnected results towards fulfillment of a particular objective. The model however built primarily from the angles of convergence of reports on possible precursor molecules, factors leading to circularization and cellular spatiotemporal condition supporting circularization.

Towards resolving the first problem of finding precursor molecule, as for primary guess, first off, each of the exonic circular RNAs collected from 2 databases were BLASTed against the lncRNAs distributed within 3 databases to see the region of homology with 100% similarity to ascertain at primary level that lncRNAs have the region inside their sequence from which their circular form may be originated in particular condition. As for supportive reports we find that circular RNA mostly originated from exonic regions. Other regions for their origin are also reported like, UTR and Intronic regions, lncRNA loci and antisense of known transcripts [24,25]. Also, Recent report reveals that ciRS-7 exonic sequence is embedded in an lncRNA locus [26]. Taking cue from these works and focusing this study on circular RNAs rooted from lncRNAs, a circular RNA query processing mechanism against lncRNA database was performed as discussed above. Subsequently the lncRNA data providing mapped hits and no hits were collected for further study. The target of this part of work was to find the similarity between lncRNAs and Circular RNAs at the level of sequence so that the precursor molecule (possibly lncRNA) of Circular RNA could be identified.

Next part of this work was primarily devoted to identify biomolecular factors and the spatiotemporal conditions catalyzing circularization of these precursor molecules (notionally lncRNAs as indicated in this study) producing Circular RNA. In this regard, RNA binding proteins (RBPs) common to both lncRNAs and Circular RNAs [27,28] were intuitively targeted as possible important biomolecular catalysts for circularization of RNAs for their reported roles in such activities [4,5,7]. The cellular spatiotemporal conditions supporting circularization of RNAs were also studied on the basis of differential profile of existence (DPE) of number of lncRNAs and Circular RNAs within nucleus and cytosol of a cell in association with DPE of RBPs unique to lncRNAs and Circular RNAs. The objective of studying spatiotemporal conditions leading to formation of Circular RNAs was to ascertain whether their biogenesis is co-transcriptional or post-transcriptional.

## Materials & methods

### Integrated Cellular Geography (ICG) formalism to study biogenesis of circular RNA

Integrated Geography (IG) in its definition is the branch of geography that describes and explains the spatial aspects of interactions between human individuals or societies and their natural environment, called coupled human–environment systems [23]. Since biological cell can be thought of as a micro-geographic space, Integrated Cellular Geography (ICG) was employed as an alternative Systems Engineering approach to get more in-depth information about the system through pulling in apparently unconnected published results and putting them together under a framework of Data Science [35, 35b]. In this study, the equivalence of IG with ICG was drawn by substituting human by concerned biomolecules and geographic environment by cellular environment. A mathematical framework of this approach is given below.

For a pool of logical lemma, 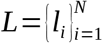, if these lemmas

i. serve as supports to prove theorem, *T* = *f* (*l*), for f is a function in broader sense which is derived through judiciously made association of all the members of L, and,
ii. represent complimentary form of necessary and sufficient proof of T,

We may consider the analysis of the system is compatible and doable under the broad paradigm of Integrated Data Geography (IDG) and specifically, for this work, Integrated Cellular Geography (ICG).

In our work which dealt with biomechanical root of formation of circular RNA, the geographical space of our choice was a biological cell and its very important sub-cellular spaces (nucleus and cytosol) from where the referred biomolecule was generated. We analyzed the spread of this biomolecular entity outside and inside the sub-cellular space along with assimilating other reported pieces of information (e.g., about RBP molecules involved in circularization of such molecules) to come up with necessary and sufficient conditions of root of formation of such molecule.

### Collection of data and Pre processing

#### Data collection strategy

The objective of data collection for this work was primarily to study the effectiveness of the newly proposed Integrated Cellular Geography approach in predicting biogenesis of circular RNA. Under this approach, following two types of data were considered for their utilization:

i. Data type 1 (DT1): Apparently unconnected reports of published papers, and,
ii. Data type 2 (DT2): Formal data collected from databases.

Therefore for the model constructed under this approach, the intention was to look into the convergence of DT1 and DT2 to extract information on:

i. possible precursor molecules of circular RNA,
ii. molecular agents involved in circularization, and,
iii. cellular spatiotemporal condition supporting circularization.

#### Collection of data for identifying possible precursor molecules of circular RNA

Considering the specific class of circular RNA that were exonic and were intuitively formed from lncRNAs, the sequences of circular RNAs were collected from circBase [36] and circRNADb [37], while lncRNAs sequence were downloaded from GENCODE [14], NONCODE [38] and LNCipedia [39]. The current version of the dataset was downloaded like gencode.v19.lncRNA_transcripts.fa from GENCODE, high confidence lncRNAs lncipedia_5_0_hc.fasta from LNCipedia and NONCODEv5_human.fa from NONCODE. These datasets were pre-processed by writing Perl scripts to extract relevant information for further study.

#### Collection of data for identifying factors leading to circularization of RNA

As mentioned in the introduction section, RNA Binding Proteins were reported to be involved in the circularization of some RNAs. To get a comprehensive view and extract relevant information, circInteractome [27], starBase [28] and POSTAR[40] databases were used.

#### Collection of data for detecting cellular spatiotemporal condition supporting circularization

For this purpose, online database and web server were used by taking specific queries like, RBP-types or lncRNAs to search out their special location inside cellular spaces. UniProtKB [41] was used for getting the annotation of RBPs which included their localization information. Also web server iLoc-LncRNA [42] was used to get locations of lncRNAs within cellular spaces.

### Identification of possible precursor molecules of circular RNA

#### Mapping the circular RNAs sequence on lncRNAs

In this phase, the objective was to search out the region of homology of circular RNAs on lncRNAs to initially check if there was any indication for lncRNAs to serve as precursor molecules of circular RNAs. For this purpose, we have taken two circular RNA databases, circBase and circRNADb in our study. Also for lncRNAs we have taken datasets from three lncRNAs database like GENCODE, LNCipedia and NONCODE. Following dataset, query-set and tools were used for the mapping:

i. Datasets for reference data of BLAST [43] tool: GENCODE, LNCipedia and NONCODE
ii. Query-set: Queries obtained from circular RNA datasets, circBase and circRNADb
iii. Search tool: BLASTN

Mapping process was given in Table 6.

### Molecular agents involved in circularization

#### Study on RBP types in relation to their binding with lncRNA and circular RNA along with differential profile of their existence within nucleus and cytosol following ICG

As clarified in the introduction section, in this work, role of some RBPs was studied in relation to circularization of RNA. Therefore, it was important to get account of their spatial existence especially within nucleus and cytosol of a cell along with information on the types of RNAs they bound. Therefore, databases, circInteractome [27], starBase [28] and POSTAR [40] were utilized to extract information on RBPs unique to Circular and lncRNAs as well as common to both. Subsequently percentage existences of RBPs for each of the RNA-types along with RBPs unique to both Circular and lncRNAs were computed in:

i. Cytosol only
ii. Nucleus only
iii. Both nucleus and cytosol, and,
iv. Secreted from the cell

To accomplish this task, supplementary files extracted from databases, CircInteractome and starBase were utilized to extract RBPs and RNAs they bound to by writing Perl scripts to read these files. However, in case of POSTAR database following methodological steps were utilized to extract list of RBPs for lncRNAs:

Step 1). First, POSTAR.csv file was downloaded and the list of lncRNAs was extracted by parsing of this file.
Step 2). Each of the lncRNAs within the list was submitted as query to POSTAR to find out the RBPs having binding sites on it.
Step 3). The RBPs extracted in step2 were further processed to eliminate redundant instances of them.

Furthermore, to get information on spatial location of RBPs, each one of them was inputted into UniProtKB [35] to get its spatial location from the output annotated data.

#### Identification of QKI and FUS RBPs, as common to both Circular and lncRNAs

Among many different types of functions mediated by QKI type of RBPs, in this work, its role for circularization was investigated from the report [5]. This report highlighted that:

i. Epithelial to mesenchymal transition (EMT) coincides with regulation of many circular RNAs,
ii. Circular RNA formation was regulated by QKI during EMT,
iii. Binding sites of QKI was found to be flanking circRNA-forming exons, and,
iv. Circularization of linear RNA could be induced by insertion of QKI binding site into them.

Similarly the role of another RBP, FUS in regulating circularization was documented in [34]. Therefore, it was imperative in this study to investigate on RBPs common to both Circular and lncRNAs which was carried out using existing literatures and databases, POSTAR and CircInteractome. POSTAR provided the information about the post transcriptional regulatory role of RBPs while CircInteractome gave information of circular RNAs, their RBPs and microRNAs. Towards this direction, the methodology adopted comprised of two stages:

Stage1: At this stage following steps were applied to get first hand information about RNAs for a specific RBP-type:

Step 1: Specific RBP (either QKI or FUS) was submitted as query in POSTAR database.
Step 2: The database returned the names of the genes and corresponding RBP-type binding sites on RNAs transcribed from these genes.

Stage2: Since direct description of RNAs (whether mRNA or lncRNA) could not be extracted at stage 1, following steps were applied to get such information:

Step 1: Out of the output gene pools extracted through stage 1, only those transcribing for lncRNAs were screened through writing Perl script.
Step 2: Each of the screened genes was submitted as query in CircInteractome database and the outputs yielding at least 1 hit were chosen only.
Step 3: Each of the outputs obtained from step 2 was further analyzed to check manually the RNA types transcribed by this gene (say, Circular RNA, since this database checks for Circular RNA only), RBP types for these RNA types (say, QKI or FUS) and binding sites for these RBPs on these RNA types. Only those genes transcribing for Circular RNA with specific RBP-types (QKI or FUS) were retained for further analysis.

### Cellular spatiotemporal condition supporting circularization

#### Exploring cellular locations of lncRNAs having sequence similarity with exonic circular RNAs

As described in subheading under the methodology section “Identification of possible precursor molecules of circular RNA”, to examine whether exonic circular RNAs can be originated from lncRNAs following steps were performed using fixed number of Circular RNAs and lncRNAs obtained from the process mentioned above.

1. Based on the total BLAST hits obtained only those hits were extracted (A) in which the circular RNAs sequences fully mapped in lncRNAs.
2. Using A, specific web-server, iLoc-LncRNA (‘http://lin-group.cn/server/iLoc-LncRNA/predictor.php’) was used to know probable spatial location of these lncRNAs inside the cellular space.

#### Exploring cellular locations of lncRNAs having no sequence similarity with exonic circular RNAs

Following steps were performed for this purpose:

1. Based on the total BLAST hits obtained only those hits were extracted (A) in which the circular RNAs sequences were not at all mapped in lncRNAs.
2. Finally cellular location of such lncRNAs was found using iLoc-LncRNA web server. To reduce computational complexity in dealing with very large database where number of data is greater than 1000, sample datasets were used through random selection of data from the original database.

Workflow depicting the ICG model for biogenesis of circular RNA was shown in Fig. 1.

**Fig. 1.**
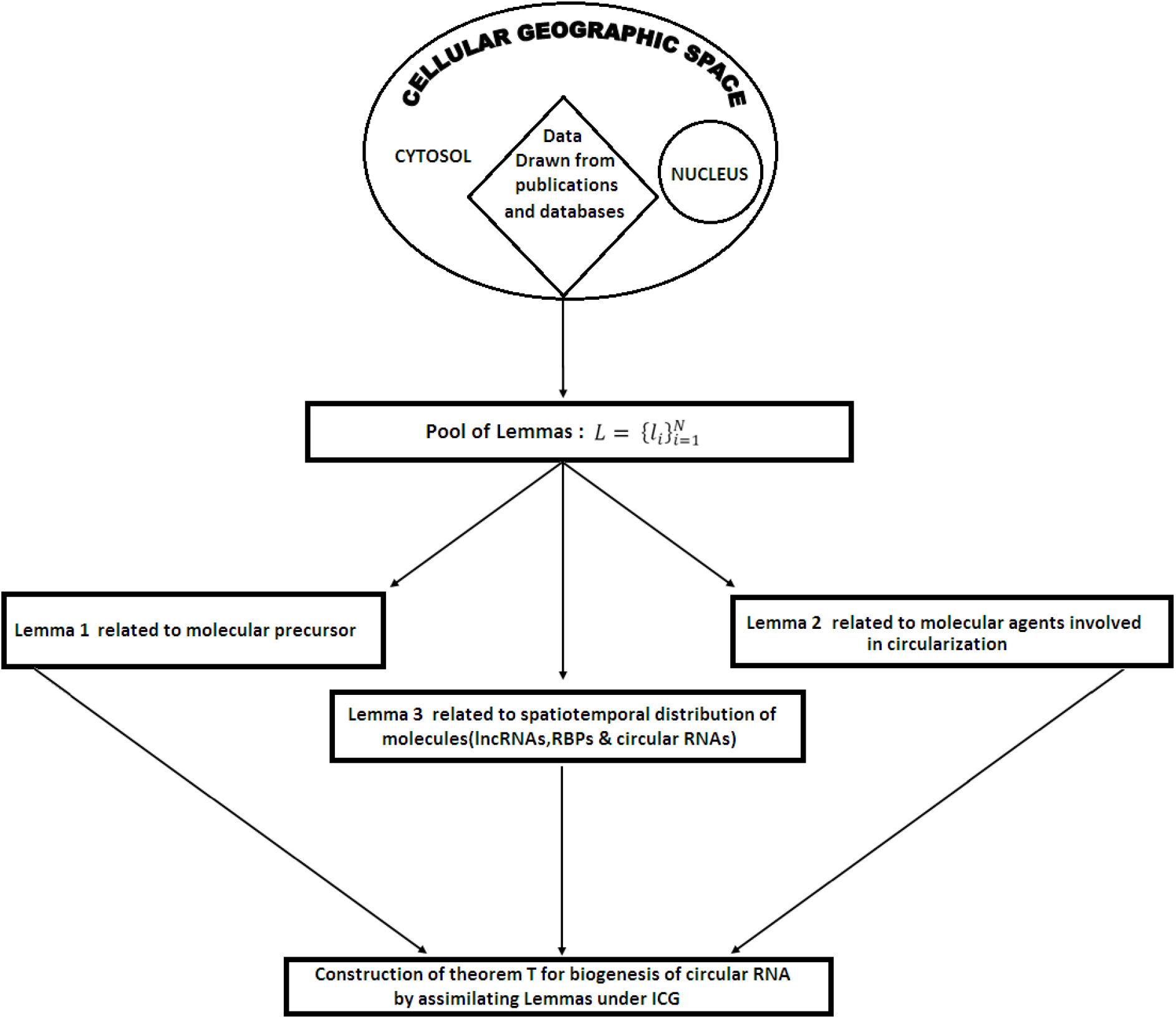
Workflow based ICG model for biogenesis of circular RNA

## Results

As stated in the introduction section, the objective of this work was to study molecular mechanism leading to formation of Circular RNA. To accomplice it, first possible precursor molecules were searched and thereafter, possible factors leading to conversion of these precursor molecules into Circular RNA were studied. Therefore, the results were listed in a manner so that each of the results lead to formation of a logical lemma that served as a building block to construct the proposed ICG based model of biogenesis. A simple chemical representation of this model was shown in Fig.2 which was made the basis for the extraction of the results.

**Fig.2.**
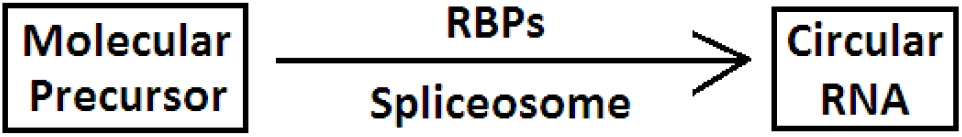
Chemical representation of ICG

### Result to construct lemma 1 related to identification of potential molecular precursors of Circular RNA

In this regard, methodologies as described in methodology section under subheading “Identification of possible precursor molecules of circular RNA” was employed to check status of lncRNAs towards such function because degeneracy of transcription from both coding genes and non-coding nucleotide sequence producing them was well reported [29]. The numbers of lncRNAs sequences in GENCODE, NONCODE and LNCipedia under study were 23898, 172216 and 104487 respectively. The total numbers of circular RNAs taken from circRNADb, circBase in study were 32914 and 140790 respectively. The total number lncRNA hits including those that were completely mapped from particular lncRNA databases against circular RNAs as queries was shown in Table 1.

**Table 1.** lncRNAs hits against circular RNAs as queries

| Circular RNA database | lncRNAs in database |  |  | BLAST HITS information |
| --- | --- | --- | --- | --- |
|  | GENCODE | NONCODE | LNCipedia |  |
| circRNADb | 1211 | 7471 | 1846 | Total hits obtained |
|  | 170 | 907 | 184 | Hits yielding complete mapping with query circular RNA |
| circBase | 7977 | 37339 | 13010 | Total hits obtained |
|  | 135 | 1620 | 205 | Hits yielding complete mapping with query circular RNA |

Lemma 1: Reasonable percentage of lncRNA (~5%) out of total hits obtained was found to have complete sequence similarity with circular RNA as shown in Table 1 indicating lncRNA as possible precursor molecule of circular RNA.

### Result to construct lemma 2 related to identification of RBPs as molecular agent for biogenesis

The lemma related to identification of RBPs as possible molecular agent stood on the following conjectures:

1. RBPs helping circularization from precursor lncRNA to circular RNA should first attach with lncRNA and remain attached till the process of circularization.
2. For the above conjecture to be true, RBPs solely binding with lncRNA or circular RNA might not be considered for molecular agent helping circularization.
3. RBPs common to lncRNA and circular RNA might be considered as agent for the circularization of RNA.
4. For the conjectures 1 to 3 to be true, there must be existence of reasonable amount of common RBPs

Furthermore, for validation of this lemma role of RBPs common to both lncRNA and circular RNA was explored. Towards this direction, methodology section under subheading “Molecular agents involved in circularization” was utilized to further corroborate this result with the help of RBPs unique to Circular and lncRNAs and common to both. Results of those RBPs were obtained from starBase, Circinteractome and POSTAR. Fig.3 showed Venn diagram generated using Venny 2.1.0 tool [30] to show binding distribution of RBPs with different RNA types. In the pool of RBPs common to both RNA types, presence of QKI and FUS was found to be important to validate the lemma since both of these RBPs were reported to be involved in circularization.

**Fig. 3.**
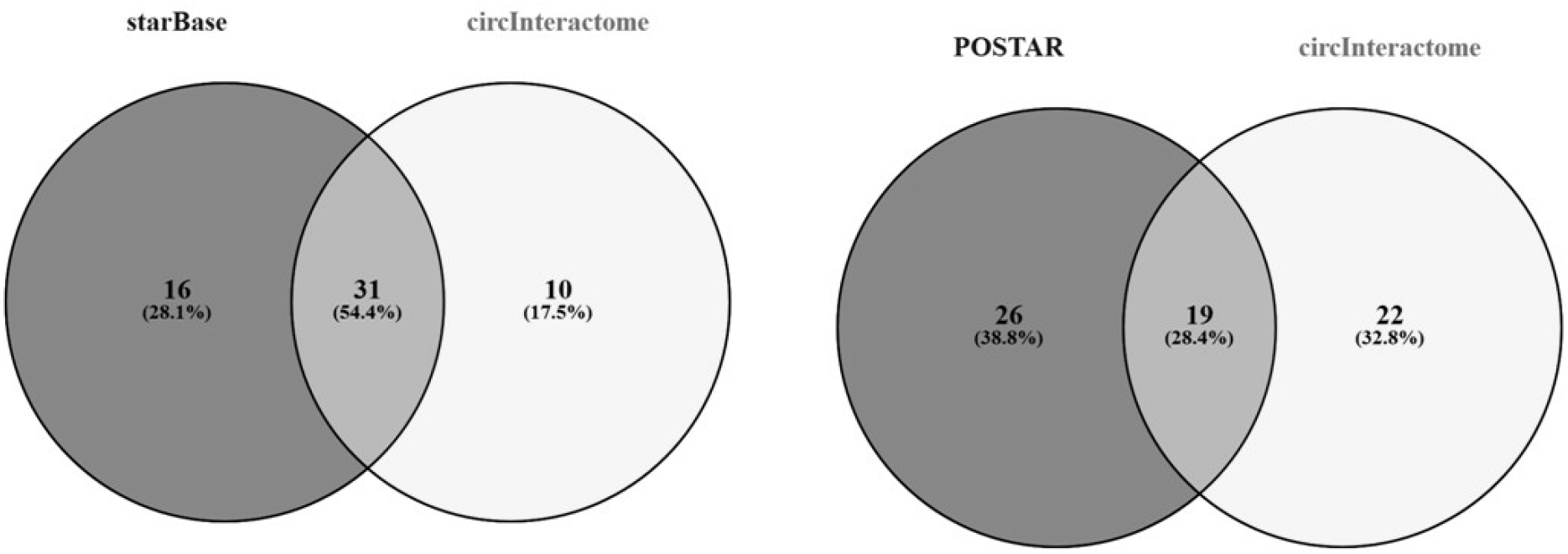
Venn diagram showing percentage of RBP-types bound to lncRNAs and Circular RNAs

Also, for validating both lemma 1 and 2, steps as described in methodology section under subheading “Identification of QKI and FUS RBPs, as common to both Circular and lncRNAs”, were used to extract information on number of genes producing both lncRNA and circular RNA transcripts whereas, both of these transcripts had binding sites for FUS or QKI as shown in Table 2.

**Table 2.** Common genes transcribing both RNA types with binding sites for FUS and QKI

| **RBP** | **lncRNA type** | **A** | **B** | **C** |
| --- | --- | --- | --- | --- |
| FUS | lincRNA | 104 | 9 | 6 |
|  | antisense | 73 | 5 | 3 |
|  | sense_intronic | 10 | 2 | 0 |
|  | sense_overlapping | 9 | 1 | 0 |
| QKI | lincRNA | 47 | 6 | 1 |
|  | antisense | 24 | 0 | 0 |
|  | sense_intronic | 1 | 1 | 0 |
|  | sense_overlapping | 2 | 0 | 0 |

[Counts of: screened genes transcribing only lncRNAs obtained as hits from POSTAR database (A), hits of genes returned against screened genes submitted as query in CircInteractome (B) and genes returned by CircInteractome transcribing RNAs having binding sites for FUS and QKI (C)]

Lemma 2: Some of the RBPs common to both test lncRNA and circular RNA served as molecular agent for circularization of lncRNA.

### Result to construct lemma 3 related to spatio-molecular distribution of lncRNA, exonic circular RNA and RBPs

Using the methodology section under subheading “Exploring cellular locations of lncRNAs having sequence similarity with exonic circular RNAs” and “Exploring cellular locations of lncRNAs having no sequence similarity with exonic circular RNAs” in Methodology section, the localization profile of lncRNA was obtained and shown in Table 3 and Table 4 respectively.

**Table 3.** Cellular spatio-molecular distribution of lncRNAs having sequence similarity with exonic circular RNA

| Query<br>Circular<br>RNA<br>database | Distribution in Cellular Space(in %) |  |  |  | Reference<br>lncRNA<br>Database |
| --- | --- | --- | --- | --- | --- |
|  | Cytoplasm | Nucleus | Ribosome | Exosome |  |
| CircRNADB | 63.529 | 23.529 | 05.294 | 07.647 | GENCODE |
|  | 59.782 | 25.543 | 08.152 | 06.521 | Lncipedia |
|  | 58.875 | 24.807 | 06.945 | 09.261 | NONCODE |
| CircBase | 58.846 | 23.589 | 10.256 | 07.307 | GENCODE |
|  | 57.900 | 24.900 | 11.700 | 05.500 | Lncipedia |
|  | 55.700 | 23.100 | 10.800 | 08.300 | NONCODE |
| Average | 59.105± 2.577 | 24.245±0.968 | 8.858± 2.476 | 7.423±1.321 |  |

**Table 4.** Cellular spatio-molecular distribution of lncRNAs which do not have any notable sequence similarity with exonic circular RNA

| Query Circular RNA database | Distribution in Cellular Space(in %) |  |  |  | Reference lncRNA Database |
| --- | --- | --- | --- | --- | --- |
|  | Cytoplasm | Nucleus | Ribosome | Exosome |  |
| CircRNADB | 52.700 | 24.000 | 12.800 | 10.500 | GENCODE |
|  | 53.000 | 25.000 | 13.800 | 08.200 | Lncipedia |
|  | 54.200 | 23.800 | 10.800 | 11.200 | NONCODE |
| CircBase | 51.500 | 25.800 | 13.700 | 09.000 | GENCODE |
|  | 51.000 | 25.900 | 13.000 | 10.100 | Lncipedia |
|  | 54.400 | 25.400 | 10.000 | 10.200 | NONCODE |
| Average | 52.8± 1.378 | 24.983± 0.899 | 12.35± 1.580 | 9.867±1.084 |  |

As described in methodology section under subheading “Study on RBP types in relation to their binding with lncRNA and circular RNA along with differential profile of their existence within nucleus and cytosol following ICG”, to obtain spatial distribution of RBPs, only those RBPs were considered, sub cellular localization of which was reported in UniProtKB. As shown in Table 5, it was evident that RBPs that specifically bound to Circular RNA types were more localized within cytosol rather than nucleus of a cell and the contrary was true in case of RBPs bound to lncRNA types.

**Table 5.** The spatial distribution profile of RBPs unique for RNA types Circular and lncRNA

| RNA type with which RBPs bound | Databases providing Information of RBPs related to RNA types | Sub cellular distribution of RBPs (in %) obtained from UniProtKB database against RBPs as queries |  |  |  |
| --- | --- | --- | --- | --- | --- |
|  |  | Cytosol only | nucleus only | Both nucleus and cytosol | Secreted |
| Circular RNA only | CircInteractome | 26.90 ± 3.10 | 19.29 ± 9.29 | 53.81 ± 6.19 | 0 ± 0 |
| lncRNA only | starBase& POSTAR | 10 ± 10 | 51.88 ± 8.13 | 28.85 ± 8.85 | 9.38 ± 9.38 |
| Both lncRNA and circular RNA | CircInteractome, starBase & POSTAR | 18.60 ± 3.48 | 22.41± 9.36 | 58.99 ± 5.88 | 0 ± 0 |

**Table 6.** Steps for mapping of circular RNA query onto lncRNA reference dataset

| Mapping Steps | Executable used | Parameter | Value of parameters |
| --- | --- | --- | --- |
| Creation of BLAST database | makeblastdb | -in | lncRNA database sequences |
|  |  | -input_type | 'fasta' |
|  |  | -dbtype | 'nucl' |
|  |  | -out | BLAST_generated_lncRNA_database_name |
| Searching homology | blastn | -query | circular RNAs database sequences |
|  |  | -task | 'blastn' |
|  |  | -db | BLAST_generated_lncRNA_database_name |
|  |  | -evalue | 1e-05 |
|  |  | -outfmt | 6 |
|  |  | -perc_identity | 100 |
|  |  | -num_threads | 8 |
|  |  | -out | Output_file name |
|  |  | -max_target_seqs | 1 |
|  |  | -max_hsps | 1 |

The process was performed for all possible combinations of query and reference databases.

Facts obtained from Table 3, 4 and 5 and publications report as building blocks of lemma 3 were as follows:

1. In general, lncRNAs were more localized within cytosol rather than nucleus of a cell.
2. Percentage of lncRNAs within cytosol that had considerable sequence similarity with circular RNA, was more than that having no notable sequence similarity with circular RNAs.
3. Exonic circular RNAs were more localized within cytosol [21,25,31] and outcome of Table 3.
4. Percentage of RBPs which bound with circular RNA only and localized both in nucleus and cytosol was much more than that localized in cytosol or nucleus only. This fact together with fact 3 indicated that these RBPs bound with minor fraction of exonic circular RNAs already residing within nucleus, and therefore can be considered to be formed within nucleus.
5. Percentage of RBPs which bound with lncRNA only and localized in nucleus was much more than that localized in cytosol or both cytosol and nucleus. This fact together with facts 1, 2 and 3 indicated that these RBPs had hardly any role in circularization.
6. Percentage of RBPs which were common both for lncRNA and circular RNA and localized in both nucleus and cytosol was much more that localized in cytosol or nucleus only. Furthermore, focusing on localization of this type of RBP in cytosol only or nucleus only, their frequency was found to be reasonably greater in nucleus than cytosol. This facts together with facts 1, 2, 3 and 4 indicated that these RBPs played more significant role in circularization of RNA in comparison to other RBP types.
7. QKI was localized both in cytosol and nucleus whereas FUS was localized in nucleus.

Lemma 3: Probability that RBPs common to both lncRNA and circular RNA first binds with lncRNA within nucleus for circularization and is transported together with lncRNA to cytosol for final circularization into exonic circular RNA form is very high along with some probability that the same RBP type lead to formation of exonic circular RNA within nucleus.

### Integrated cellular geography approach to elucidate biogenesis of Circular RNA

As stated earlier in introduction, methodology and initial part of result section, ICG approach considered to build a theorem upon the bases of lemmas supporting it. Following this approach we first built the proposition in the form of a model that there should be existence of a precursor molecule of circular RNA which upon the involvement of some molecular agents would yield exonic circular RNA as shown in Fig.1.

Towards this direction, the lemmas (lemma 1, 2 and 3) as a result of outcomes extracted in this work from various published experiments (as shown in Tables 1 to 5) led to formation of following theorem:

#### Theorem on biogenesis of exonic circular RNA

Biogenesis of exonic circular RNA is mostly post-transcriptional event which starts with the binding of RBPs common to both lncRNAs and circular RNAs with precursor molecule lncRNAs within nucleus and ends within cytosol with some exceptions following spatio-molecular arrangements of lncRNA, circular RNA and all types of RBPs.

## Discussion

In this work, mechanism of biogenesis of exonic circular RNA was studied from the angle of sequence semantics and assimilation of apparently unconnected published data under the paradigm of ICG as described earlier. Sequence semantics helped in identification of precursor molecule as lncRNA that was supposed to convert into circular RNA form. The biogenesis mechanism was studied considering the spatio-molecular aspect of lncRNAs, circular RNAs and RBPs within cellular space. Overall, ICG was employed to decipher the mechanism of biogenesis of circular RNA under the theoretical formalism as described in methodology section under subheading “Integrated Cellular Geography (ICG) formalism to study biogenesis of circular RNA”. The concept of ICG was pressed in for such task as a generalization of naturally evolved solution instead of being forced upon into this study. The reason was a plethora of already published reports on experiments done on circular RNA and allied biomolecules which were although unconnected apparently in their present form, yet could be studied further in their assimilated form for confirming a target objective, i.e., a theorem. Although methodology for assimilation of published data, like Meta Analysis existed, it was found to be applicable only for data generated using same experimental protocol. The essence of ICG was actually that of Integrated Geography at microscopic, i.e., cellular scale which was further generalized in this work in a logical lemma based formalism as especially shown in the methodology section under subheading “Integrated Cellular Geography (ICG) formalism to study biogenesis of circular RNA” and the result section.

Circular RNAs are known as biomarkers of quite a number of diseases including cancers. This work was motivated from this fact to partially contribute to the knowledge on events leading to biogenesis of Circular RNA assuming it to be central to unearth pathobiochemical mechanism of the these diseases. In this regard, the problem was divided into two segments, first to search for molecular precursors being converted into circular form and identification of factors leading to its circularization to produce Circular RNA. Secondly, the objective was to extract mechanism of biogenesis of circular RNA from the spatio-molecular distribution of lncRNA, circular RNA and RBPs. As described in the result section, lemmas were extracted as supports for solving both of these problems. These lemmas were constructed using existing databases and results of published works as research materials and through judicious assimilation of those under ICG.

To solve the first problem of identifying molecular precursor for Circular RNAs, lncRNAs were considered as primary guess material through such indication from the published research works [15,26]. Since there was no solid evidence or proof so far for lncRNAs being circularized to form Circular RNAs, semantic similarity in sequence order between lncRNAs and Circular RNAs was first explored to establish physicospatial relationship between these two entities. Moreover, the lncRNAs following the rules set in this study were found to have resemblance with antisense properties, which was one more attribute common for Circular RNA [21,24,32]. A good proportion (~32%) of antisense transcripts in human was reported as lncRNAs [14]. These two findings indicated fulfillment of necessary condition for lncRNAs to be considered as molecular precursor of Circular RNA. In an apparent contradiction, although lncRNAs were sometimes found to be transcribed from coding genes, it was intriguing to find its ultimate form as whole or partial natural antisense (i.e., non-coding) transcripts (NAT) as reported by Milligan & Lipovich et al. (2015) [33]. To check authenticity of this primary guess apart from fulfilling the requisite necessary conditions, methodology section described in subheading “Identification of possible precursor molecules of circular RNA” was employed in this work which searched for existence of completely mapped lncRNA sequence-hits against circular RNA sequence as query. Table 1 of result section under subheading “Result to construct lemma 1 related to identification of potential molecular precursors of Circular RNA” provided the outcome of this exercise from which the lemma 1 was extracted. According to lemma 1, “reasonable percentage of lncRNA (~5%) out of total hits obtained was found to have complete sequence similarity with circular RNA as shown in Table 1 indicating lncRNA as possible precursor molecule of circular RNA”.

In continuation with first problem, identification of factors leading to circularization of RNA was tried to be resolved using methodology section under subheading “Molecular agents involved in circularization” the result of which was shown in result section under subheading “Result to construct lemma 2 related to identification of RBPs as molecular agent for biogenesis”. In this regard, the conjectures of result section under subheading “Result to construct lemma 2 related to identification of RBPs as molecular agent for biogenesis” were built to hypothesize about the possible class of RBPs responsible for biogenesis of circular RNA through its circularization. In this regard, Fig.3 showed the binding distribution of RBPs with different RNA types indicating presence of quite high amount of RBPs (~40.32% on an average) common to both lncRNAs and circular RNAs. Authenticity of this result was further validated through the study on the RBPs, FUS and QKI which were known as responsible for biogenesis of circular RNA and also common to both lncRNAs and circular RNAs as described in methodology section under subheading “Identification of QKI and FUS RBPs, as common to both Circular and lncRNAs” and outcome shown in Table 2. All these results together with the conjectures provided the basis of the construction of lemma 2 which was: “Some of the RBPs common to both test lncRNA and circular RNA served as molecular agent for circularization of lncRNA”. It was intriguing to find that lemma 2 stood as a support for lemma 1 also.

Regarding solution of the second problem, in this work it appeared to be interesting to extract spatio-molecular root of Circular RNA via its relationship with RBPs and earlier molecular form i.e., as indicated from this study, lncRNAs. That said, problem reduced to find out overall spatial distribution of lncRNAs, Circular RNAs and RBPs within nucleus and cytosol of cells. Regarding this, the results shown in Table, 3, 4 and 5 using methodology sections, under subheading “Exploring cellular locations of lncRNAs having sequence similarity with exonic circular RNAs”, subheading “Exploring cellular locations of lncRNAs having no sequence similarity with exonic circular RNAs” and subheading “Study on RBP types in relation to their binding with lncRNA and circular RNA along with differential profile of their existence within nucleus and cytosol following ICG” together with the facts generated in result under subheading “Result to construct lemma 3 related to spatio-molecular distribution of lncRNA, exonic circular RNA and RBPs” provided the basis for formulation of lemma 3 which was: “Probability that RBPs common to both lncRNA and circular RNA first binds with lncRNA within nucleus for circularization and is transported together with lncRNA to cytosol for final circularization into exonic circular RNA form is very high along with some probability that the same RBP type lead to formation of exonic circular RNA within nucleus”.

In the context of utilization of existing databases and apparently unconnected published reports, as stated in the beginning of this discussion and methodology section under subheading “Integrated Cellular Geography (ICG) formalism to study biogenesis of circular RNA”, it appears to be quite justified to employ the formalism of integrated cellular geography (ICG) following the concept of general Integrated Geography (IG) [23]. It grossly described spatial nature of relationship between objects, the environment holding them and the world containing both of them, which found their equivalence with RBPs, RNA types and whole cellular space comprising of cytosol and nucleus respectively. The main purpose of employing this approach was to find the origin and historical dynamism of an object in relation to the spatio-molecular environment holding it. Interestingly the lemmas extracted out of this approach appeared to be complementarily supportive to each other to lead to the final theorem on biogenesis of exonic circular RNA which was: “Biogenesis of exonic circular RNA is mostly post-transcriptional event which starts with the binding of RBPs common to both lncRNAs and circular RNAs with precursor molecule lncRNAs within nucleus and ends within cytosol with some exceptions following spatio-molecular arrangements of lncRNAs, circular RNAs and all types of RBPs”. The salient feature of this approach is that, similar to lemma based study of a mathematical theorem, complexity in reasoning leading to an inference from apparently unconnected biological data and reports can be reduced substantially where lemmas served as filtered knowledge.

The interesting part of this work was to consider studying the role of RBPs in circularization of RNAs. In this regard, it appeared to be quite obvious that RBPs specific for Circular RNAs might not have any role for their biogenesis since they would bind with circular form only, which was not possible before circularization. The inference stemmed from the same reasoning led to the fact that RBPs common to both of these RNA types should be prime molecular factors that could be expected to help circularization of lncRNAs and thus would have to remain attached throughout transition of lncRNA to Circular RNA. This fact was further supported by the presence of such RBPs, QKI and FUS with their established role in circularization [5,34]. However, in this work the role of QKI and FUS as a common RBP of both Circular and lncRNA types could also be found from analyses described in methodology section under subheading “Study on RBP types in relation to their binding with lncRNA and circular RNA along with differential profile of their existence within nucleus and cytosol following ICG” and Table 5 of result section under subheading “Result to construct lemma 3 related to spatio-molecular distribution of lncRNA, exonic circular RNA and RBPs”. Therefore, role of other common RBPs remained to be an interesting area of research in relation to their role as molecular factor for circularization. Fig.4 further clarified this model. Furthermore, as reported by Barrett et al. (2017), a Circular RNA, ciRS-7 was also found to be embedded in LINC00632 which was an lncRNA. This fact was also found to be in support of our theorem [26].

**Fig4.**
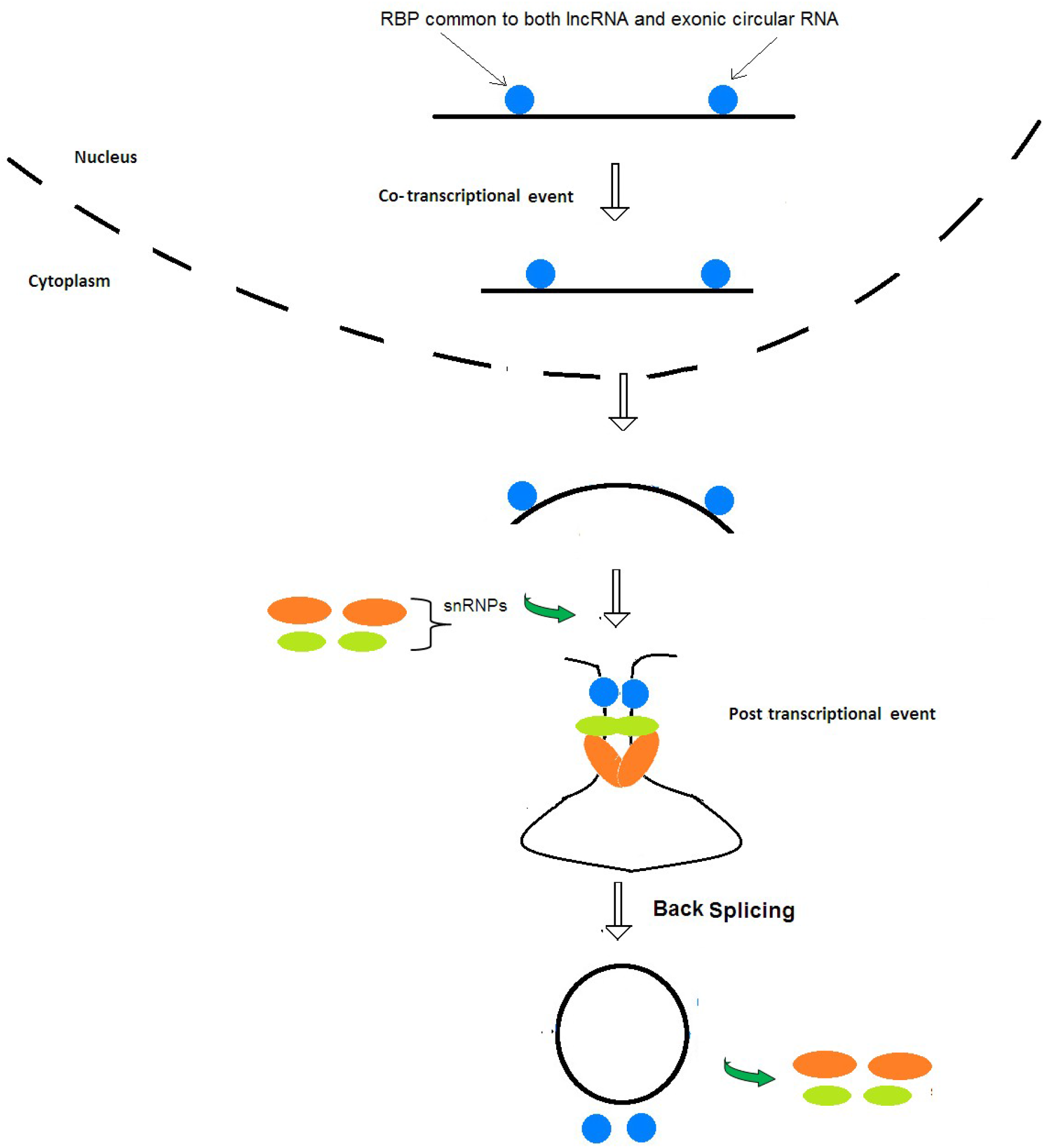
Model describing different stages of circularization of lncRNA (black line and curves) with the help of RBPs common to both Circular (black circles) and lncRNAs.

However, for RBPs specific for lncRNAs the chance of their having role in the transition from linear to circular form of RNA being less following the above-mentioned reasoning, discussion on them was avoided although possibility of their involvement might not be overruled.

## Conclusion

Overall, this study showed a possible model of biogenesis of exonic circular RNAs from lncRNAs as precursor molecule on the foundation of judicious fusion of results and data of already performed experiments under the paradigm of integrated cellular geography approach as analytical basis. Sequence analysis of both RNA types revealed significant similarity making the foundation of a logical lemma based study related to possibility of biogenesis of Circular RNAs from lncRNAs. Additionally, information extracted in this work from distribution of these RNA types within cellular spaces, cytosol and nucleus, in relation to their binding with specific types of RBPs further helped in building of other two lemmas in support of spatio-molecular mechanism of biogenesis of exonic circular RNA. The theorem built from these three complimentarily supportive lemmas strongly supported model of biogenesis of Circular RNA as provided in this work. According to this model: biogenesis of exonic circular RNA begins within nucleus with the binding of RBPs common to both lncRNAs and circular RNAs with precursor molecule lncRNAs and ends within cytosol with some exceptions following spatio-molecular arrangements of lncRNAs, circular RNAs and all types of RBPs. Therefore, this model indicated that the biogenesis of exonic circular RNA was mostly a post-transcriptional event. The concept of ICG, as first time introduced in this work, appeared to have prospective of a very useful methodological ingredient of systems biology for holistic study of events of cellular system.

## Acknowledgements

We are thankful to Department of Applied Science, IIIT-Allahabad, India for allowing uninterrupted use of computer systems. Rajnish Kumar and Manoj Kumar Pal are grateful to IIIT-Allahabad for providing them fellowship to pursue their doctoral work.

## Availability of data and materials

This study utilizes the data and web services which are already available in public domain.

## Authors’ contributions

Rajnish highlighted the importance of study of circular RNA from which Tapobrata understood importance of exploring biogenesis of circular RNA and convinced other authors about it. Rajnish proposed and worked towards finding the molecular precursor of circular RNA through analysis of sequence semantics which has been accepted by other authors. Tapobrata proposed study of cellular environment and role of proteins towards circularization of the precursor molecules. Rajnish introduced all about the possible interesting roles of RNA Binding Proteins (RBPs) in circularization. In this regard, Tapobrata strategized the cellular environmental study of circular RNA, lncRNA and RBPS incorporating already published experimental results under the lemma-based framework of Integrated Cellular Geography model and Rajnish implemented it. Author, Manoj was with all these activities both in conceptualization and implementation part.

## Ethics approval and consent to participate

Not applicable.

## Consent for publication

Not applicable.

